# MolMap resolves functional chromatin responses to small-molecule perturbation

**DOI:** 10.64898/2026.11.16.766766

**Authors:** Eliska Fürst, Eva Karina Brinkman, Adriana Carvajal-Jimenez, Mengjun Wu, Vicent Pelechano, Ilaria Piazza

**Author notes:** Shared corresponding authors.

## Abstract

Small molecules can alter chromatin through protein-, DNA- and physicochemical interactions, yet current approaches do not directly resolve where these perturbations produce an immediate functional chromatin response in native nuclear environments. Here we present MolMap (*chromatin-small Molecule Mapping*), a sequencing-based method that profiles genome-wide chromatin responses to chemically unmodified compounds in isolated, non-crosslinked nuclei.

Rather than measuring ligand binding, protein occupancy or downstream gene expression, MolMap identifies genomic regions where small-molecule exposure produces an immediate functional chromatin response. By quantifying changes in Tn5 accessibility and local chromatin organization in native nuclei, MolMap captures locus-specific and dose-dependent effects while separating early chromatin responses from downstream cellular adaptation. Across metabolites, protein-targeting compounds, a DNA-binding drug and a physicochemical chromatin modulator, MolMap resolved compound-specific and mechanistically distinct responses. Comparison with live-cell ATAC-seq showed that MolMap captures early chromatin responses before broader cellular adaptation. In leukemia models, MolMap revealed a coordinated chromatin response in drug-sensitive cells that was strongly attenuated in resistant models. MolMap therefore provides a native framework for bridging the precision of biochemical perturbation assays with the breadth of genome-wide profiling, extending classical chromatin biochemistry into a genomic setting.

## Main

In eukaryotic cells, genetic information is packaged in chromatin, a dynamic assembly of DNA and associated histone and non-histone proteins. Chromatin governs gene expression through the action of protein complexes that regulate epigenetic processes, including histone or nucleic acid modifications and chromatin remodeling. RNA-binding regulators, architectural proteins, histone modifying enzymes and transcription factors (TFs) collectively shape chromatin structure and function, enabling context-specific regulation of gene activity ^1^.

Because chromatin architecture is central to cell identity and gene regulation, it is highly responsive to small molecules that act through chromatin-associated proteins and DNA structures. Many of these processes are also therapeutically tractable^2^, and several classes of chromatin-targeting compounds— including inhibitors of histone deacetylases (HDACs) and DNA or histone methyltransferases—are clinically established or under active development ^3–5^. Despite this, determining where small-molecule exposure produces an immediate chromatin response within a native nuclear environment remains challenging ^5–8^.

Existing approaches interrogate different aspects of small-molecule–chromatin interactions. *In vitro* biochemical assays can quantify ligand binding or enzymatic activity ^9–11^, but generally lack native chromatin context and genome-wide resolution. Occupancy-based methods can localize small-molecule binding across the genome in cells but require fixation and compound derivatization, introducing potential technical artifacts and limiting throughput ^12–14^. Likewise, proximity labelling approaches can identify protein interaction partners on chromatin but may require gene editing ^15,16^, whereas biophysical methods such as fluorescence recovery after photobleaching ^17,18^ report binding dynamics but not locus-resolved responses. Chromatin proteomics provides complementary information on chromatin-associated protein composition ^19,20^, while transcriptomic and proteomic profiling measure downstream cellular output. The broader transcriptional and regulatory programs can mask the initial molecular events that follow compound treatment, complicating the identification of the molecular processes most closely associated with ligand action. Resolving such proximal effects can provide mechanistic information relevant to compound optimization, ligand analog design and the identification of potential *off-target* effects.

Collectively, current approaches capture complementary aspects of small-molecule action on chromatin: binding, occupancy, dynamics, composition and downstream cellular responses, but none directly measure the immediate chromatin response to small-molecule perturbation in native nuclei. There is therefore a critical need for genome-wide tools that detect proximal chromatin responses to small molecules in their native biochemical context, before they become intertwined with secondary cellular adaptation.

To address this gap, we developed **MolMap** (*chromatin-small Molecule Mapping*), a method that profiles early chromatin responses to chemical perturbations. By using changes in chromatin structure and local organization as a readout of compound activity, MolMap reveals proximal molecular consequences of small-molecule ligand engagement at genome-wide scale. Because compounds are applied directly to native nuclei outside of living cells, MolMap isolates chromatin-level responses from downstream gene expression and signalling changes. This enables unbiased, quantitative interrogation of how small-molecule ligands perturb chromatin and provides insight into their mode of action within a native nuclear context.

## Results

### MolMap captures proximal chromatin responses to chemical perturbations

#### MolMap

(*chromatin-small Molecule Mapping*) enables genome-wide profiling of chromatin responses triggered by small-molecule perturbation, independently of downstream cellular effects. The assay is performed in native, non-crosslinked nuclei extracted from cells, allowing for precise control of compound concentrations and exposure times. Compounds are directly added to these nuclei, where active gene expression and signaling have been halted, allowing immediate biochemical chromatin responses to be measured without downstream transcriptional or signalling feedback. Compound-induced chromatin responses are then quantified by measuring genome-wide changes in Tn5 transposase accessibility (**Figure 1a**). MolMap therefore uses changes in accessibility and local chromatin organization to resolve proximal chromatin responses to chemical perturbation.

**Figure 1.**
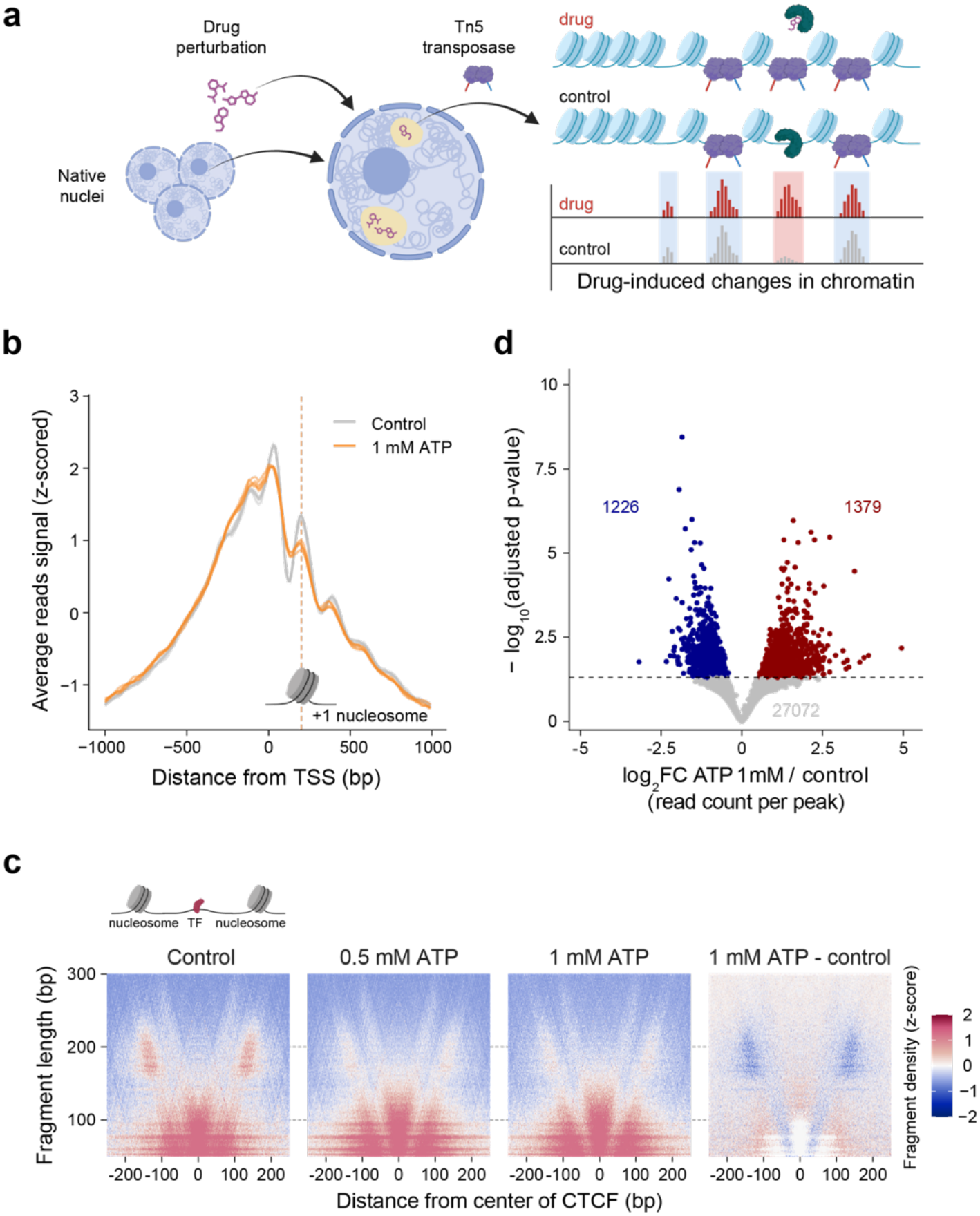
MolMap captures proximal chromatin responses to chemical perturbations. **a**. The MolMap workflow. Native nuclei are isolated from cells and exposed to small molecules under biochemical conditions that preserve chromatin structure while halting cellular responses within a time frame of minutes. Tn5 transposase fragments accessible chromatin regions and the resulting DNA fragments are sequenced to quantify compound-induced chromatin changes. **b**. Average MolMap signal profiles centered on transcription start sites (TSSs) ±1 kb, following treatment with standard buffer or 1 mM ATP. Signals represent z-scored, library size–normalized coverage per 10 bp bin, averaged across 28,278 TSSs (refGene annotation; n = 4 biological replicates). Dashed lines indicate the +1 nucleosome region, where ATP treatment alters the accessibility profile. **c**. Fragment length vs. position heatmaps (V-plots) showing z-score normalized MolMap fragment density around CTCF associated sites at each position and size with intensity indicating read enrichment in K562 nuclei under control, 0.5 mM ATP, or 1 mM ATP. The x-axis denotes the distance (±250 bp) from the center of binding sites, and the y-axis represents the length of MolMap fragments (in base pairs). The central short-fragment protection pattern corresponds to the CTCF-associated site; flanking patterns reflect nucleosome positioning. The final panel shows the difference between 1 mM ATP and control. **d**. Volcano plot showing ATP-induced chromatin structural changes. Each point represents a MolMap peak; the x-axis shows log_2_ fold change (1 mM ATP vs. control) and the y-axis shows –log_10_ adjusted p-value (DESeq2 Wald test, adjusted p < 0.05; Benjamini–Hochberg correction). Peaks with significant increased or decreased accessibility are shown in red and blue, respectively.

We define *chromatin structural changes* as both sharp accessibility shifts (e.g., peak changes) and more subtle alterations in Tn5 tagmentation patterns, such as changes in nucleosome organization or protein-associated protection patterns ^21^. Whereas conventional ATAC-Seq ^22^ measures chromatin accessibility in a cellular context, MolMap applies defined perturbations directly to isolated nuclei over short incubation times. This experimental design separates the immediate chromatin response to a defined chemical perturbation from secondary cellular responses, providing a genome-wide biochemical readout of where ligand exposure produces measurable changes in chromatin structure and local organization *in situ*. We operationally define loci exhibiting such changes as functionally responsive chromatin regions.

To benchmark the method, we first tested the effect of ATP, a naturally occurring metabolite that fuels ATP-dependent chromatin remodelers and modulates nucleosome organization ^23,24^. Addition of ATP to native nuclei at physiological concentrations (0.5 and 1 mM) altered the characteristic phased nucleosome accessibility pattern around transcription start sites (TSSs)^25^, with a pronounced change at the +1 nucleosome region. This response is consistent with ATP-dependent chromatin remodelling^26^ (**Figure 1b and Extended Data Fig. 1a**). This effect was abolished when ATP was replaced with the non-hydrolysable analogue AMPPNP, indicating that the observed chromatin changes reflect immediate functional consequences of ligand engagement, rather than ligand binding alone (**Extended Data Fig. 1b)**. ATP also caused dose-dependent structural changes at CTCF, MYC, and PHF8 -associated sites (**Figures 1c and Extended Data Fig. 1c**), affecting both transcription factor-occupied and nucleosome-protected regions.

MolMap therefore does not directly measure protein occupancy or enzymatic activity, but instead captures chromatin structural changes that arise as immediate functional consequences of ligand engagement. MolMap provides a mechanism-agnostic readout of chromatin response, irrespective of whether the underlying mechanism involves altered protein binding, catalytic activity, chromatin remodeling or other regulatory mechanisms.

Across the genome, MolMap detected significant differential accessibility for 2,605 peaks (adjusted p-value < 0.05) in response to 1 mM ATP (**Figure 1d**), and for 90 peaks with 0.5 mM ATP. Notably, 993 of the ATP-responsive peaks at 1 mM overlapped with TSSs (accounting for 38% of all changes), highlighting ATP’s direct influence on nucleosome organization near 5’ of actively transcribed genes ^23^. These results demonstrate that MolMap captures metabolite-induced chromatin structural changes with high sensitivity and resolution.

To assess the robustness of the assay, we tested an alternative buffer condition with increased ionic strength ^27^. ATP-induced chromatin structural changes were preserved under these conditions, with comparable MolMap differential accessibility profiles observed between the standard and higher-stringency MolMap buffers (**Figure 1b and Extended Data Fig. 1d**). These results indicate that MolMap readouts are robust to variations in ionic conditions, and we proceeded with the standard buffer for subsequent experiments.

### MolMap resolves chromatin responses to protein-targeting drugs

Having estabilished that MolMap can resolve genome-wide chromatin responses to a metabolite in native nuclei, we next applied the method to protein-targeting drugs. We selected four chemically diverse compounds with distinct mechanisms of action: topotecan (topoisomerase I inhibitor), etoposide (topoisomerase II inhibitor), GSK3368715 (type I protein arginine methyltransferase inhibitor), and trichostatin A (a histone deacetylase inhibitor).

MolMap revealed compound- and mechanism-dependent chromatin-response signatures that reflected the distinct modes of action of each drug. Principal component analysis (PCA) of peak accessibility showed strong clustering by treatment condition, with the first two components explaining 59% and 15% of the variance, respectively (**Figure 2a**). While ATP induced widespread chromatin accessibility changes, protein-targeting drugs produced more site-specific effects. We therefore focused on accessibility changes at defined regulatory elements to test MolMap’s sensitivity to subtle perturbations.

**Figure 2:**
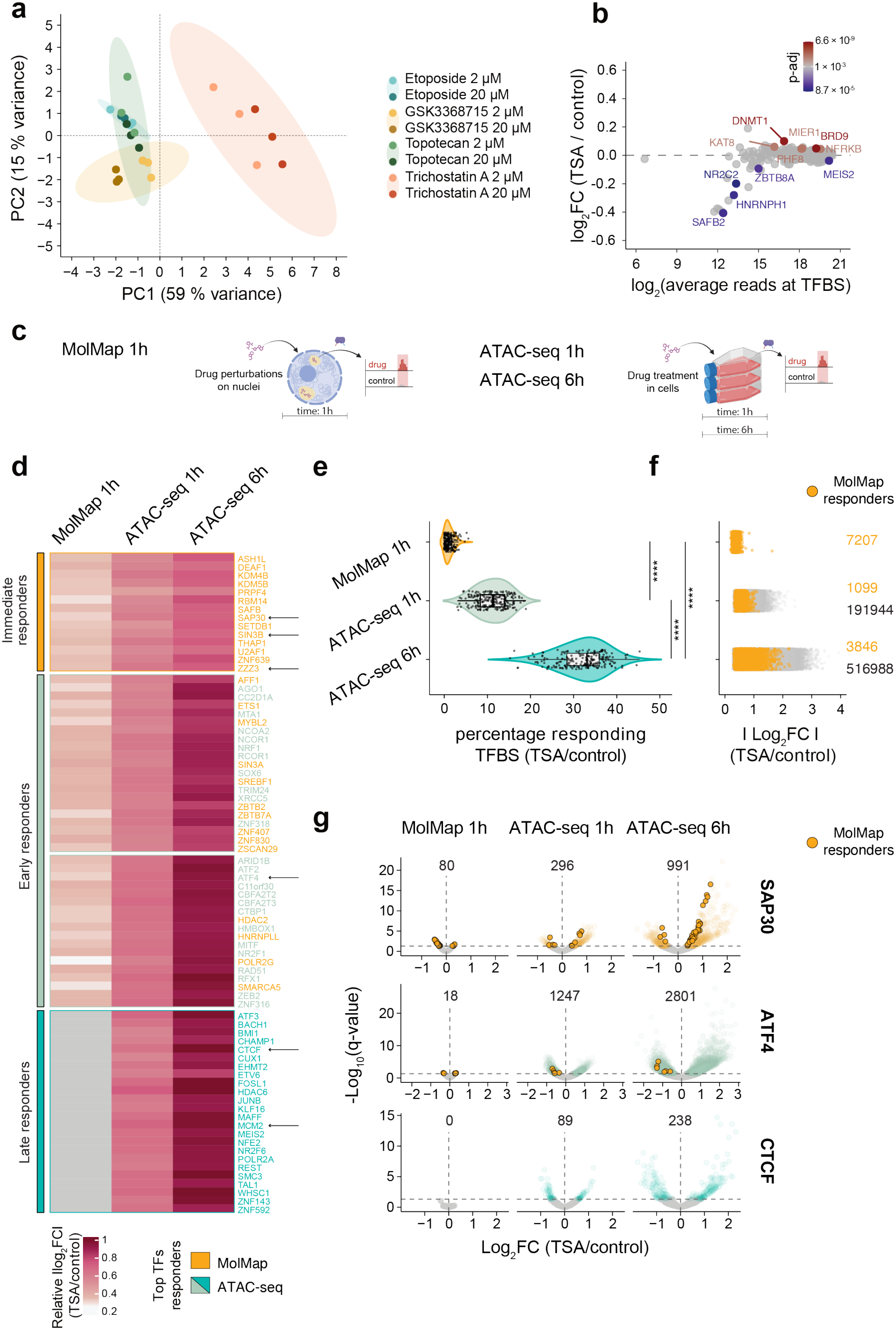
Dissecting biochemical and cellular chromatin responses with MolMap and ATAC-seq. **a**. Principal component analysis (PCA) of compound-induced MolMap chromatin accessibility changes across drug treatments. Each point represents an individual biological replicate. Colored areas show 95% confidence intervals grouped by drug treatment. **b**. MA plots of differential MolMap accessibility at aggregated transcription factor-associated sites following treatment of K562 nuclei with 20 µM of trichostatin A. Each point represents one TF and their color indicates the direction and significance of accessibility changes (DESeq2 Wald test; adjusted p < 0.001, Benjamini–Hochberg correction), with red and blue showing increased and decreased accessibility. **c**. Experimental set-up of the time matched MolMap and ATAC-seq assays with 20 µM of Trichostatin A (TSA). Isolated nuclei were subjected to TSA treatment for 1 hour (MolMap 1h). Live cells were treated with TSA for 1 or 6 hours (ATAC-seq 1h, ATAC-seq 6h). **d**. Heat map of chromatin accessibility changes at transcription factors-associated sites (TFBS) upon 20 µM of Trichostatin-A treatment in MolMap 1h, ATAC-seq 1h and ATAC-seq 6h assays. Each row shows the top 30 ranked TF responses grouped in 4 clusters after separation by unsupervised k-mean clustering. For each TFs the responses are computed as the median of absolute values of log_2_ fold changes for all significantly changing TFBS in in MolMap 1h, ATAC-seq 1h and ATAC-seq 6h (DESeq2 Wald test, adjusted p < 0.05, Benjamini–Hochberg correction). Min = 0.25, Max = 1.04. **e**. Distribution of percentages of transcription factor-associated sites (TFBS) significantly changing accessibility for all mapped TFs in MolMap 1h, ATAC-seq 1h and ATAC-seq 6h after treatment with 20 µM trichostatin-A (TSA). Each point indicates a TF (DESeq2 Wald test, adjusted p < 0.05, Benjamini– Hochberg correction; violin plot: horizontal lines define the median and boxes the 25th and 75th percentiles; whiskers: 1.5 × the interquartile range). **** indicate statistically significant changes of the median of the three groups (Wilcoxon test, adjusted p < 0,0001). **f**. Differential TFBS accessibility changes upon 20 µM of Trichostatin A (TSA) treatment in MolMap 1h, ATAC-seq 1h and ATAC-seq 6h (numbers on top indicate the total number of TFBS significantly changing in the three assays; DESeq2 Wald test; adjusted p < 0.05, Benjamini–Hochberg correction). Each point indicates an individual TFBS with responses computed as the absolute value of log_2_ fold changes. Orange points pinpoint TFBSs with a significant change in MolMap signal. **g**. Volcano plots of accessibility changes at individual chromatin regulators sites (SAP30, ATF4 and CTCF) in MolMap 1h, ATAC-seq 1h and ATAC-seq 6h upon 20 µM of Trichostatin A (TSA) treatment. Each point represents a TFBS. Orange points pinpoint TFBSs with a significant change in MolMap signal; they are in large part also significantly changing loci in the ATAC-seq datasets. Numbers above each plot indicate the number of significant TFBSs detected in the corresponding dataset (DESeq2 Wald test, adjusted p < 0.05, Benjamini–Hochberg correction).

Topotecan and etoposide, which both inhibit topoisomerases, elicited comparable early chromatin responses. Topotecan altered accessibility at 27 transcription factor (TF)- associated sites (adjusted p < 0.001; **Extended Data Fig. 2a**), and etoposide affected 44 of them (adjusted p < 0.001; **Extended Data Fig. 2b**). Both compounds preferentially impacted loci associated with RNA polymerase II (POLR2G) and spliceosome-related factors (for instance, RBFOX2 and PYGO2; **Extended Data Fig. 2a-c**) or DNA damage-related proteins (XRCC5, BRCA1). These effects are consistent with the role of topoisomerases in relieving torsional stress during transcription elongation and RNA splicing ^28^. Inhibiting their activity disrupts supercoiling dynamics, potentially impairing co-transcriptional processing.

GSK3368715 is a type I PRMT inhibitor, and likely exerts its effects mainly through inhibition of PRMT1 ^29^, the primary enzyme responsible for asymmetric di-methylation of H4R3 ^30^ and subsequent euchromatin formation with chromatin opening ^31–33^. Consistent with this, GSK3368715 reduced accessibility at TSSs (**Extended Data Fig. 2d**). Upon PRMT1 inhibition, chromatin accessibility changed for instance at HDAC2- and KDM5B-associated sites (**Extended Data Fig. 2e**), indicating that these regions are particularly sensitive to PRMT1-dependent chromatin regulation.

We next examined how MolMap captures chromatin responses to epigenetic regulators that directly modulate histone modifications. Trichostatin A treatment increased chromatin accessibility at multiple transcription factor– bound regions, including HDAC-associated repression complexes (MIER1), chromatin remodelling (NFRKB, BRD9) and chromatin acetylation machinery (KAT8) (adjusted p < 0.001; **Figure 2b**). This pattern is consistent with trichostatin A’s function as a histone deacetylase inhibitor, which promotes H3K27 hyperacetylation and enhances chromatin accessibility. Accordingly, we observed a global increase in accessibility at annotated H3K27ac regions (**Extended Data Fig. 2f**).

To directly link MolMap accessibility changes to specific genomics sites, we implemented two antibody-guided tagmentation approaches in isolated native nuclei under MolMap conditions: a Tn5-based targeted MolMap (analogous to CUT&Tag) and a MNase-based targeted MolMap (analogous to CUT&RUN) (**Extended Data Fig. 2g**). The Tn5-based targeted MolMap readout using an H3K27ac-specific antibody confirmed that trichostatin A–induced accessibility changes co-localize with increased H3K27ac at defined genomic loci (**Extended Data Fig. 2h**).

### Dissecting direct and indirect chromatin responses across biochemical and cellular contexts

Having established that MolMap captures immediate chromatin structural responses to histone acetylation, we next asked how these primary events compare to chromatin accessibility changes measured in living cells, where transcriptional feedback and secondary signaling are expected to progressively amplify chromatin responses over time, potentially obscuring the direct targets of drug action. To this end, we compared MolMap with matched ATAC-seq experiments following trichostatin A treatment in isolated nuclei and intact cells (**Figure 2c**).

We grouped TF-associated sites by their accessibility-response profiles across MolMap and ATAC-seq over time. This unsupervised, data-driven clustering separated early MolMap-enriched responders from factors that became progressively responsive in cells over time upon trichostatin-A treatment in 4 clusters (**Figure 2d**). The MolMap-enriched cluster (**Figure 2d, top cluster**) contained loci associated with canonical HDAC repression complexes, including SAP30 and SIN3B, as well as ZZZ3, a component of the ATAC histone acetyltransferase complex, consistent with direct perturbation of the acetylation– deacetylation balance following HDAC inhibition (**Figure 2d, top cluster**). Thus, MolMap preferentially captured chromatin regions linked to the primary biochemical axis of trichostatin A activity. These factors represent direct chromatin-acetylation regulators and immediate responders to TSA exposure.

In contrast, ATAC-seq in living cells revealed a progressive expansion in both the magnitude and number of accessibility changes over time, increasing from hundreds of TF-associated sites at 1 h to thousands at 6 h after treatment (**Figures 2e and 2f**). Early ATAC-seq responses partially overlapped with MolMap (**Figure 2d, middle clusters**), but also included additional transcription factors associated with rapid enhancer activation, such as ATF4, which are weakly or not detected in MolMap (**Figure 2d, middle clusters**; **Figure 2g**). At later time points, accessibility changes extended to higher-order chromatin regulators (CTCF, MCM2, **Figure 2d, bottom cluster; Figure 2g**) and to stress-responsive factors, reflecting widespread transcription-coupled remodeling.

TFs identified by MolMap showed limited changes in effect sizes across time points in ATAC-seq, whereas many ATAC-seq–specific factors showed progressively increasing accessibility changes (**Figure 2f and Figure 2g, Extended Data Fig. 2i**). This supports a model in which MolMap preferentially captures primary chromatin regulatory nodes engaged by the compound, while ATAC-seq reflects downstream amplification of these effects through transcriptional and signaling networks.

Together, these findings establish a temporal and mechanistic distinction between the two approaches: MolMap captures primary, locus-specific chromatin structural responses to compound engagement, whereas ATAC-seq reflects progressively amplified, system-level chromatin changes driven by cellular adaptation.

These findings highlight MolMap’s ability to detect both broad and localized chromatin changes in response to protein-targeting drugs. By quantitatively resolving compound-specific alterations in chromatin accessibility, MolMap enables mechanistic dissection of drug action directly within native chromatin.

### Polyamine-induced chromatin compaction is resolved by MolMap in native nuclei

To evaluate whether MolMap can capture global chromatin structural changes driven by intrinsic biophysical properties of chromatin, we applied the assay to isolated nuclei exposed to spermidine, a physiological polyamine known to promote chromatin compaction through electrostatic charge neutralization, independent of sequence-specific DNA recognition or specific protein-ligand binding ^34^. Consistent with a global shift toward a more compact chromatin state, spermidine treatment resulted in an approximately two-fold reduction in normalized library yield, indicative of decreased genome-wide accessibility (**Figure 3a**). At the same time, transcription start site (TSS) profiles became markedly sharpened, with narrower and more pronounced accessibility peaks relative to control conditions (**Figure 3b**).

**Figure 3:**
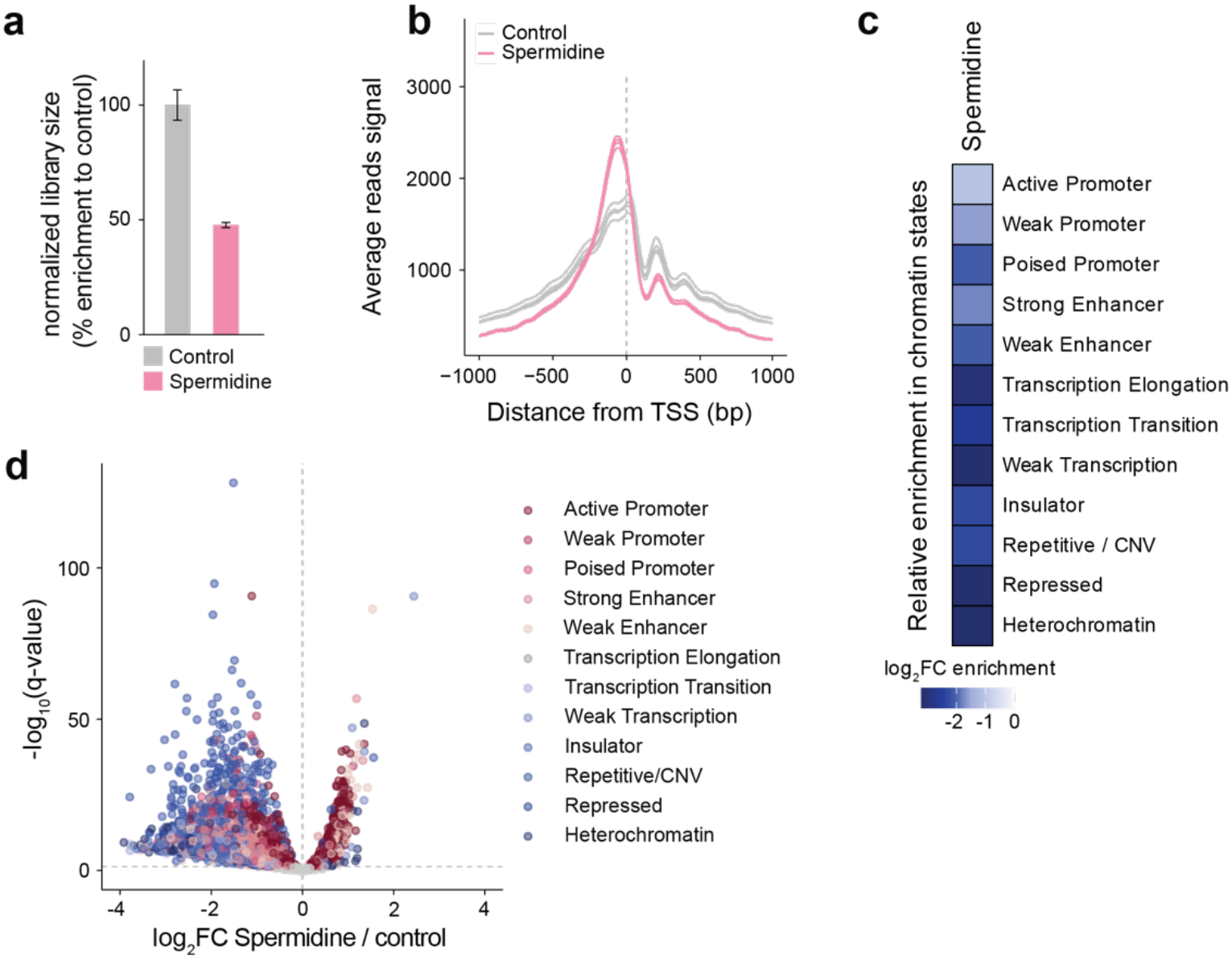
Polyamine-induced chromatin compaction is resolved by MolMap in native nuclei. **a**. Relative MolMap library size following 5 mM spermidine treatment. Library sizes were normalized to the internal *E. coli* DNA reference and scaled to the average DMSO control (100%). Error bars represent standard deviations to the mean (n ≥ 4). **b**. Average MolMap signal profiles at transcription start sites (TSSs) ±1 kb for control (DMSO) and 5 mM spermidine-treated nuclei. Signals represent normalized read coverage per 10 bp bin, averaged across all TSSs. **c**. Heatmap of chromatin state-specific MolMap accessibility changes following 5 mM spermidine treatment. Values represent log_2_ fold changes relative to DMSO control across ChromHMM-defined chromatin states. Normalized read counts were used. CNV=copy number variation. **d**. Volcano plot of Spermidine-induced chromatin MolMap accessibility changes visualized as log_2_ fold change (FC) between spermidine 5 mM and vehicle. Each point represents a MolMap peak. (DESeq2 Wald test, adjusted p < 0.05; Benjamini–Hochberg correction). Peaks are colored by their affiliation to a chromatin state (ChromHMM).

Because spermidine induces a global reduction in Tn5 accessibility, we use this decrease as a proxy for chromatin compaction. In the following, we refer to this as chromatin compaction level, operationally defined as the extent of accessibility reduction measured by MolMap under defined biochemical conditions.

Despite this global reduction in accessibility, differential analysis revealed a non-uniform response across the genome. Regions associated with transcriptionally inactive chromatin exhibited the strongest decreases in accessibility, whereas active regions were comparatively buffered (**Figure 3c**). Stratification by chromatin state confirmed that transcriptionally inactive regions were most affected, while gene-rich and transcriptionally active regions showed more modest changes (**Figure 3d, and Extended Data Fig. 3a**).

Chromatin compaction level was globally associated with GC content, with low-GC regions showing stronger accessibility loss (**Extended Data Fig. 3b-c**). However, GC composition alone did not explain differences between chromatin states (**Extended Data Fig. 3d**). Several states deviated from this global trend. For example, heterochromatic regions showed intermediate compaction despite low GC content (**Extended Data Fig. 3a and 3d**), while promoter states displayed distinct behaviors: active promoters were relatively resistant to compaction, whereas poised promoters were substantially more sensitive **(Extended Data Fig. 3e**), despite having comparable GC composition **(Extended Data Fig. 3d**).

Notably, heterochromatic regions that deviated from expected compaction behavior were located significantly closer to boundaries with neighboring chromatin states, suggesting that local chromatin-state transitions constrain further chromatin collapse (**Extended Data Fig. 3f**).

These findings indicate that chromatin compaction reflects not only sequence composition but also regulatory state and local chromatin organization.

The observed effects are consistent with a model in which polyamines promote chromatin compaction by bridging negatively charged DNA and nucleosomes, thereby increasing local interaction density and reducing accessibility. In this context, the preferential sensitivity of transcriptionally inactive regions suggests that chromatin domains with higher intrinsic compaction propensity are more responsive to electrostatic perturbation, consistent with previous observations in reconstituted nucleosome models ^35^.

Extending these findings to native chromatin, MolMap captures structured, locus-dependent accessibility changes arising from intrinsic biophysical properties of chromatin, rather than solely from protein-mediated regulatory events. This demonstrates that MolMap resolves chromatin responses driven by physicochemical perturbations, broadening its applicability beyond canonical ligand–protein interactions.

### MolMap reveals genome-wide chromatin disruption by a DNA-binding drug

To evaluate MolMap’s ability to capture chromatin responses to DNA-binding compounds, we profiled doxorubicin, an anthracycline that intercalates into DNA and inhibits topoisomerase II. Principal component analysis of MolMap profiles showed distinct clustering for two doses of doxorubicin, with PC1 and PC2 explaining 70% and 28% of the variance, respectively (**Figure 4a**). This shows that MolMap detects dose-dependent structural responses of a DNA-interacting compound.

**Figure 4:**
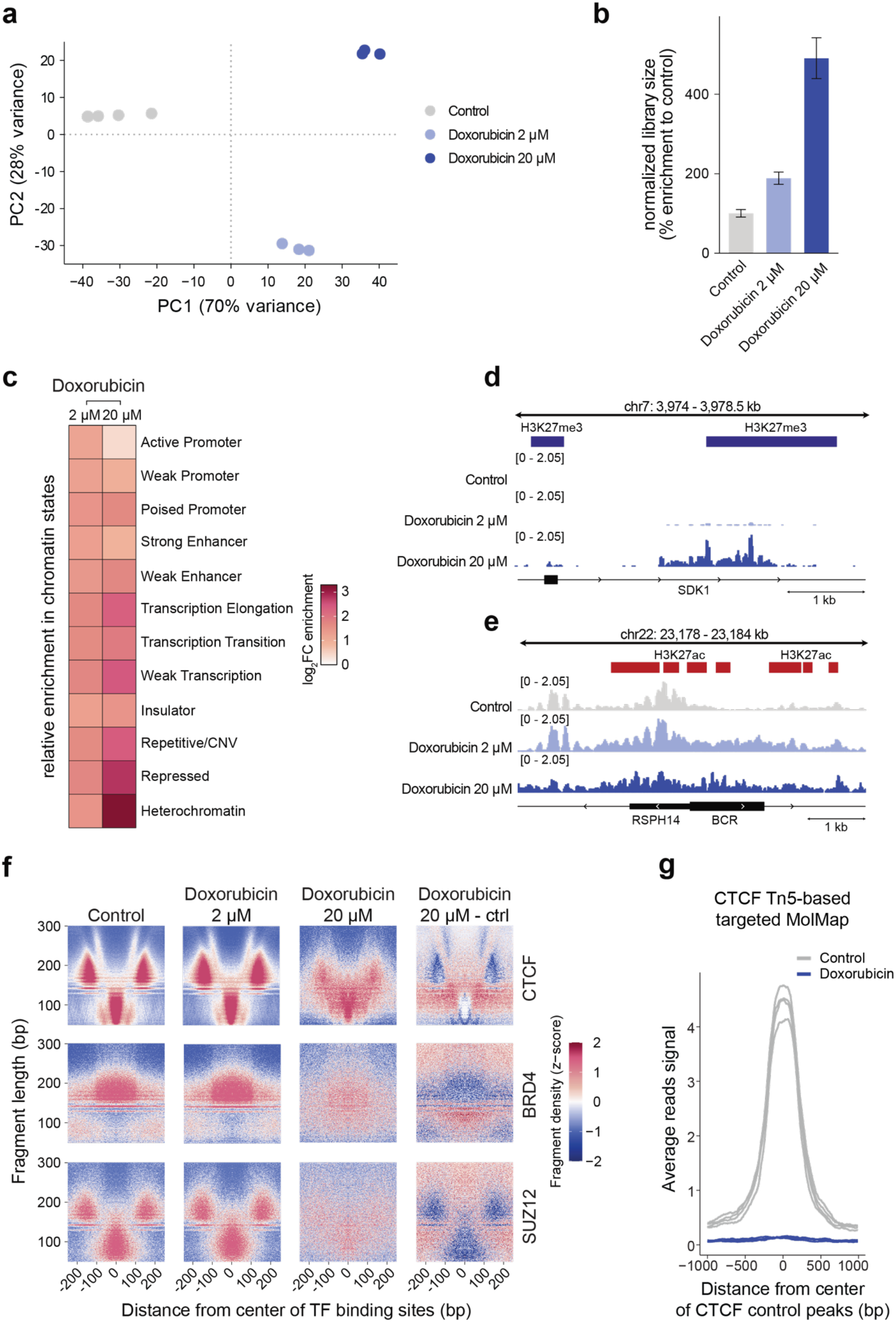
MolMap profiles genome-wide chromatin responses to a DNA-intercalating drug. **a**. Principal component analysis (PCA) of MolMap chromatin accessibility profiles after 20-minute treatment with doxorubicin (n ≥ 3). MolMap peak counts were normalized to the internal *E. coli* DNA reference. **b**. Relative MolMap library size following doxorubicin treatment analyzed using independent replicates (n ≥ 3). Library sizes were normalized to the internal E. coli DNA reference and scaled to the average DMSO control (100%). Error bars represent standard deviations to the mean. **c**. Heatmap of chromatin state-specific MolMap accessibility changes following doxorubicin treatment. Values represent log_2_ fold changes relative to DMSO control across ChromHMM-defined chromatin states. Normalized read counts were used. CNV=copy number variation. **d**. Genome browser tracks of MolMap signal at repressed (H3K27me3 containing) regions. **e**. Genome browser tracks of MolMap signal at active (H3K27ac-marked) regions. Signal tracks are from averages of three independent replicates normalized to the internal E. coli DNA reference. **f**. Fragment length versus position heatmaps (V-plots) centered on CTCF, BRD4 and SUZ12 -associated sites following doxorubicin treatment. MolMap fragment density is z-score normalized. The final panel shows the differential signal between 20 µM doxorubicin and control. Loss of short protected fragments at binding-site centers, together with altered flanking mononucleosome-sized fragments, indicates local disruption of transcription factor–associated chromatin protection and neighboring nucleosome organization. **g**. CTCF Tn5-based targeted MolMap signal profiles centered on control CTCF peaks (±1 kb) in DMSO- and doxorubicin-treated nuclei (20 µM). Signals represent CPM-normalized read coverage averaged across all peaks within the indicated genomic window relative to peak centers (n ≥ 3). Doxorubicin treatment reduces CTCF-associated chromatin signal at CTCF sites.

Doxorubicin induced a striking, dose-dependent increase in global chromatin accessibility, as quantified by normalized library size (**Figure 4b**). Accessibility gains were especially pronounced in heterochromatin and repressed chromatin states, with nine- and seven-fold enrichment, respectively (**Figure 4c**). These changes were also evident at specific genomic loci. At H3K27me3 containing regions, MolMap signal revealed strong chromatin opening in response to doxorubicin (**Figure 4d**). At active promoters enriched for H3K27ac, the core signal remained stable, whereas surrounding regions showed expanded accessibility and altered nucleosome-associated signal (**Figure 4e and 4f**). This pattern was confirmed genome-wide at transcription start sites (**Extended Data Fig. 4a**). This pattern is consistent with local disruption of nucleosome organization and is in line with previous reports that doxorubicin can destabilize histone–DNA interactions in open chromatin regions^36,37^.

Beyond nucleosome-associated changes, MolMap also detected altered protection patterns at DNA-binding protein-associated sites. V-plots centered on CTCF, BRD4 and SUZ12 -associated sites showed a dose-dependent loss of short protected fragments at motif centers, together with changes in flanking mononucleosome-sized fragments (**Figure 4f**). We interpret these profiles as local disruption of transcription factor– associated chromatin protection and neighboring nucleosome organization upon doxorubicin treatment. To validate this interpretation directly, we performed Tn5-based targeted MolMap (**Extended Data Fig. 2g**) using a CTCF-specific antibody. This confirmed reduced CTCF-associated chromatin signal after doxorubicin treatment (**Figure 4g**). Together, these data indicate that doxorubicin rapidly perturbs local chromatin architecture at defined regulatory regions, affecting both transcription factor-associated protection and adjacent nucleosome organization, likely through direct intercalation into double stranded DNA.

We next asked whether MolMap responses across the different small-molecule perturbations represented distinguishable chromatin-response patterns, rather than a generic accessibility response. To address this independently of potential sequence-dependent Tn5 insertion preferences, we derived Tn5-bias-corrected motif-associated protection scores using TOBIAS^38^, which quantify treatment-induced changes in local Tn5 protection around transcription-factor motifs across matched perturbations. Mechanistically related topoisomerase inhibitors topotecan and etoposide showed highly concordant scores, whereas GSK3368715 showed intermediate similarity and doxorubicin produced a clearly distinct response (**Extended Data Fig. 4b-c**). Consistent with the aggregated TF-level analysis (**Extended Data Fig. 2c**), topotecan and etoposide also showed concordant differential protection scores across shared TF motifs (**Extended Data Fig. 4c**). Importantly, these relative relationships were preserved after Tn5-bias correction, indicating that the observed compound-dependent response patterns are not explained by sequence-dependent transposase bias (**Extended Data Fig. 4d**). Together, these analyses show that MolMap captures structured, compound- and mechanism-dependent chromatin responses, rather than a generic consequence of exposure to small-molecule compounds.

### MolMap distinguishes BRD4-linked chromatin responses in BET inhibitor-sensitive and -resistant leukemia contexts

Identifying molecular determinants of drug resistance remains a major clinical challenge. In many cancers, resistance reflects rewiring of chromatin regulatory dependencies that are not captured by steady-state expression profiles. Nascent-transcription approaches such as SLAM-seq ^39^ quantify primary transcriptional output, while longer-term expression profiling can reveal subsequent compensatory programs ^40 41^, but they do not directly report whether drug engagement triggers structural reorganisation of protein– chromatin assemblies. MolMap interrogates a distinct molecular layer by measuring whether drug perturbation is accompanied by an immediate chromatin accessibility response in isolated nuclei. BRD4 (bromodomain-containing protein 4), a chromatin-associated transcriptional regulator of the bromodomain and extra-terminal (BET) protein family, is a major target of BET inhibition. We therefore asked whether BET inhibitor-sensitive and - resistant leukaemia contexts differ in their chromatin response to BRD4 perturbation.

To test this, we compared the MolMap response to the BET bromodomain inhibitor JQ1 ^9^ in MOLM-13 (JQ1-sensitive) and K562 (JQ1-resistant) leukaemia cells ^42^. In MOLM-13 nuclei, JQ1 induced significant accessibility changes across 22 TF-associated sites (12 increased, 10 decreased; adjusted p < 0.01; **Figure 5a**), including sites associated with BRD4 and several regulators linked to transcriptional elongation and enhancer function, including DOT1L, MED1, NELFE, CDK9/P-TEFb and SUPT5H ^43 44,45^. In contrast, under identical MolMap conditions K562 showed no significant TF-level response (**Figure 5b**). Peak-level analysis showed a similar distinction, with a broad genome-wide response in MOLM-13 and a markedly attenuated response in K562 (**Extended Data Figure 5a–b**).

**Figure 5:**
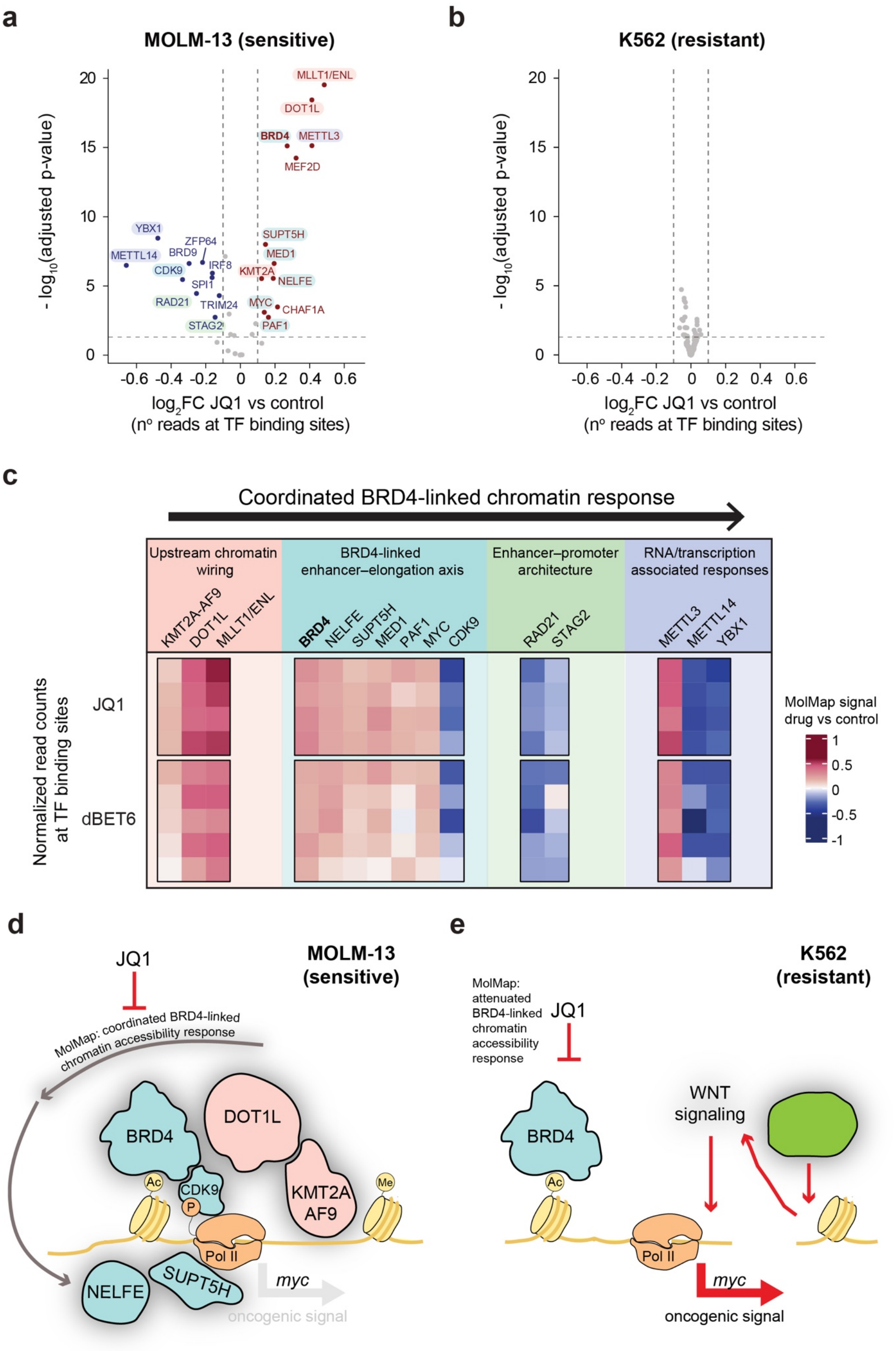
MolMap distinguishes sensitive and resistant chromatin responses to BRD4 inhibitors in leukemia cells. **a**. Volcano plot showing MolMap chromatin accessibility changes at aggregated transcription factor (TF) associated-sites in MOLM-13 cells treated with 20 µM JQ1 vs. DMSO. Red: increased accessibility; blue: decreased (DESeq2 Wald test, adjusted p < 0.01, |log_2_FC| > 0.1). **b**. Equivalent analysis in K562 cells, showing no significant chromatin response. **c**. Heatmap of selected TF-level MolMap responses following JQ1 treatment (20 µM). Normalized counts from individual control, dBET6 and JQ1 replicates are scaled and grouped according to previously described functional associations with BRD4-linked regulatory processes, including KMT2A-AF9/DOT1L regulation, enhancer–elongation, cohesin-associated enhancer–promoter organization, and RNA/transcriptional regulation. Grouping is used to aid interpretation of the coordinated BRD4-linked chromatin response. **d**. Model summarizing the coordinated BRD4-linked chromatin accessibility response detected by MolMap in MOLM-13 cells after JQ1 treatment. In this cellular context, BRD4 is functionally embedded within the KMT2A-AF9/DOT1L- and enhancer–elongation-associated regulatory machinery that supports oncogenic transcription. Upon JQ1 treatment, BRD4 inhibition is accompanied by coordinated chromatin changes across BRD4-associated and functionally linked regulatory sites, consistent with disruption of this BRD4-linked regulatory state and attenuation of the oncogenic transcriptional program. ***e***. Model summarizing the attenuated BRD4-linked MolMap response in resistant K562 cells. In this context, JQ1 does not produce the coordinated BRD4-linked MolMap chromatin accessibility response observed in MOLM-13 cells. Instead, previously described compensatory regulatory mechanisms, including WNT-dependent restoration of MYC transcription can sustain an oncogenic signal despite BET inhibition. Red arrows indicate this compensatory regulatory route described by Rathert *et al*.

Thus, MolMap distinguishes BET inhibitor-sensitive and -resistant contexts at the level of the immediate chromatin structural response, through the presence or absence of a coordinated BRD4-linked accessibility response.

Notably, this distinction is not explained simply by BRD4 engagement or by the presence of an early transcriptional response. BRD4 is present and pharmacologically targetable by JQ1 in both MOLM-13 and K562 cells, but MolMap shows that equivalent target engagement does not necessarily produce an equivalent chromatin consequence. Previous work showed that BRD4 occupancy itself poorly predicts BET inhibitors -responsive transcription, and SLAM-seq measurements demonstrated that JQ1 can elicit primary transcriptional responses of comparable scale and involving overlapping hypersensitive targets in BET inhibitor-sensitive and - resistant leukaemia models ^39^. Thus, target engagement, chromatin structural response, transcriptional output and long-term cellular sensitivity represent distinct and potentially uncoupled layers of BET inhibitor action. MolMap specifically interrogates the chromatin structural layer by asking whether BRD4 perturbation is accompanied by coordinated accessibility changes across BRD4-linked regulatory sites.

We next examined more closely the organization of the MOLM-13 MolMap response. JQ1 altered accessibility not only at BRD4-associated loci (including MYC binding-sites) but also at binding sites of factors functionally or physically linked to enhancer–promoter regulation and transcriptional elongation, including cohesin components RAD21 and STAG2 and RNA/transcription-associated factors METTL3 and METTL14 (**Figure 5a and 5c**). We refer to this pattern descriptively as a coordinated BRD4-linked chromatin response reflecting correlated accessibility changes across regulatory-factor-associated sites following BRD4 perturbation (**Figure 5c)**. This terminology does not imply that MolMap identifies the molecular pathway connecting these factors or directly measures molecular allostery.

dBET6, a BET degrader containing a JQ1-derived BRD4-binding moiety, produced a TF-level MolMap response highly similar to that induced by JQ1 (**Extended Data Fig. 5c and 5d**), indicating reproducibility across two pharmacologically related BRD4-directed perturbations.

This differential response in the two cellular contexts is consistent with known differences in the regulatory chromatin architecture of these leukaemia models. MOLM-13 cells harbour the KMT2A-AF9 fusion, which is linked to DOT1L-dependent H3K79 methylation and BRD4-associated transcriptional elongation machinery, including CDK9/P-TEFb, NELFE and SUPT5H ^46 47^ (**Figure 5d**). K562 cells lack the KMT2A-AF9 fusion protein and rely on a different regulatory architecture. Previous work^40^ has shown that JQ1 resistance in K562 reflects a defined compensatory mechanism: following the initial BRD4-dependent repression of MYC, WNT-dependent activation of a pre-existing PVT1 enhancer restores MYC transcription, whereas this bypass is absent in sensitive MOLM-13 cells (**Figure 5e**). This established regulatory rewiring provides a mechanistic context for the distinct MolMap responses observed here, without implying that MolMap itself identifies the causal resistance mechanism.

To independently examine BRD4-associated chromatin after pharmacological inhibition, we performed MNase-based BRD4-targeted MolMap in MOLM-13 cells. JQ1 treatment redistributed BRD4-associated chromatin signal rather than causing uniform loss: many loci showed reduced signal, whereas a distinct subset showed increased signal (**Extended Data Fig. 5e)**. This orthogonal assay supports the conclusion that BRD4-associated chromatin is altered following JQ1 treatment in MOLM-13, and is consistent with the coordinated chromatin response detected by untargeted MolMap.

## Discussion

MolMap provides a native, genome-wide readout of the chromatin structural consequences of small-molecule engagement, bridging controlled biochemical perturbation with locus-resolved chromatin profiling. By measuring accessibility changes in isolated nuclei, MolMap captures immediate structural responses before downstream transcriptional feedback and adaptive signaling reshape the chromatin landscape. This readout is distinct from RNA-based approaches, including SLAM-seq^39^, which can resolve immediate transcriptional output but do not directly measure the chromatin structural response that precedes or accompanies it. MolMap therefore identifies where ligand engagement is translated into a functional chromatin effect. Importantly, MolMap is target-agnostic and does not require compound derivatization, making it compatible with metabolites, DNA-binding agents and protein-targeting small molecules.

The examples in this study illustrate a common theme: immediate chromatin responses to ligand engagement are often distinct from downstream cellular outputs and are therefore missed by standard profiling assays. MolMap captures primary responders in nuclei upon treatment with HDAC inhibitor trichostatin A, such as components of the histone deacetylases and acetyltransferases complex, which are progressively masked by broader enhancer and transcriptional responses in living cells. MolMap also distinguishes the coordinated response of BRD4-associated factors upon JQ1 inhibition in JQ1-sensitive leukemia contexts. Similarly, when comparing ATP with the non-hydrolysable analogue AMPPNP, the ATP-associated change at the +1 nucleosome region was lost with AMPPNP, indicating that mere occupancy of ATP-binding sites alone is insufficient to produce the ATP-dependent chromatin response; MolMap identifies the loci where ligand exposure is accompanied by a measurable chromatin accessibility response. Together, these case studies show that MolMap identifies where ligand engagement becomes a functional chromatin response.

MolMap also captures chromatin responses driven by intrinsic biophysical properties of chromatin.

As a proof of principle case we use a net positively charged polyamine metabolite, which globally interacts with chromatin through electrostatic charge neutralization. In isolated nuclei, spermidine induced a global compaction phenotype that was nevertheless non-uniform across the genome: transcriptionally inactive and heterochromatic regions were most sensitive, whereas active promoters were comparatively buffered and boundary-proximal sites were less prone to further compaction. This indicates that chromatin state, local organization and sequence context together shape the response to a purely physicochemical perturbation. Thus, MolMap extends its applicability beyond drug responses mediated by specific ligand-binding sites to resolve ligand-induced changes in chromatin biophysics *in situ*.

Across the different perturbation classes examined here, including protein- and DNA-targeting small molecules, motif-associated differential footprinting revealed structured compound-dependent signatures, with related perturbations showing greater concordance than mechanistically distinct compounds. This supports the conclusion that MolMap does not report a generic accessibility response to small molecule exposure, but resolves where defined perturbations produce distinct chromatin structural consequences.

In leukemia models, the BRD4 experiments illustrate how MolMap interrogates a molecular layer that is distinct from both target occupancy and transcriptional output. Previous work showed that BRD4 occupancy itself is a poor predictor of BETi-sensitive transcriptional targets, and SLAM-seq demonstrated that BET inhibition can elicit primary transcriptional responses of comparable scale in BET inihibition -sensitive and -resistant leukemia models ^39^. Thus, BRD4 engagement, immediate transcriptional response and long-term cellular sensitivity are not interchangeable readouts. MolMap adds a complementary dimension by measuring whether BRD4 perturbation is accompanied by coordinated changes in the surrounding chromatin regulatory landscape.

In JQ1-sensitive MOLM-13 cells, MolMap detects coordinated accessibility changes across BRD4-associated regulatory sites and multiple factors linked to enhancer function and transcriptional elongation, whereas this response is strongly attenuated in resistant K562 cells. Importantly, the biological basis of BET-inhibitor resistance in these models has been independently dissected ^40^. Rathert et al. showed that JQ1 initially represses MYC in both sensitive and resistant leukemia cells, but resistant K562 cells rapidly restore MYC transcription through WNT-dependent activation of a distinct regulatory landscape that includes a PVT1 enhancer. Functional perturbation of this pathway alters JQ1 sensitivity in both K562 and MOLM-13, establishing regulatory rewiring rather than failure of BRD4 engagement as a determinant of resistance.

The MolMap results are consistent with this established context but report a different aspect of the response: whether BRD4 perturbation produces an immediate structural/accessibility change across BRD4-linked chromatin sites. The absence of a comparable MolMap signature in K562 therefore does not imply absence of BRD4 engagement or transcriptional responsiveness; rather, it shows that these molecular layers can be uncoupled. Importantly, previous work showed that neither steady-state transcriptomes nor acute transcriptional responses to JQ1 readily distinguish BETi-sensitive from -resistant leukemia contexts^39^. Rathert et al. were able to resolve the basis of resistance only through subsequent mechanistic dissection of enhancer rewiring and WNT-dependent restoration of MYC transcription. By contrast, MolMap reveals a distinct chromatin-response signature in MOLM-13 and K562 without prior knowledge of that compensatory pathway. This suggests a potential application of MolMap for stratifying chromatin-targeting drug responses and, when integrated with existing mechanistic or multi-omic information, for generating hypotheses about the regulatory level at which resistance emerges. More generally, this example illustrates why ligand binding, chromatin structural response, transcriptional output and eventual cellular phenotype should be considered distinct, experimentally separable layers of perturbation response.

A key limitation of MolMap is that it operates in isolated nuclei and therefore does not capture ligand responses that depend on intact cellular signalling, metabolism or cell-surface communication. However, this is also an advantage: by removing secondary feedback and intercellular variability, MolMap isolates the primary chromatin response to ligand engagement. MolMap is intentionally mechanism-agnostic: it is designed to detect where a ligand produces an immediate chromatin structural response without requiring prior assumptions about whether that response arises from altered protein binding, catalytic activity, chromatin remodeling or another regulatory mechanism. Resolving the precise molecular mechanism underlying a given MolMap response requires orthogonal follow-up experiments or complementary information; MolMap is instead intended as an unbiased readout of where a defined biochemical perturbation produces a chromatin structural response. In this sense, the assay is designed to report on proximal chromatin consequences rather than full cellular phenotypes, making it complementary to transcriptomic and proteomic profiling rather than a replacement for them.

More broadly, MolMap provides a target-agnostic framework for interrogating how defined molecular perturbations reshape chromatin in native nuclei. Its compatibility with metabolites, protein-targeting compounds, DNA-binding agents and physicochemical modulators such as sperimidine, without compound derivatization or prior knowledge of the molecular target, makes it broadly applicable to questions in chromatin biology, chemical biology and functional genomics.

By linking controlled biochemical perturbation to genome-wide chromatin readouts, MolMap extends classical chromatin biochemistry into a genomic framework.

## Supporting information

Fuerst_BioRxiv_2026_Extended

## Acknowledgements

We thank all group members of the Piazza and Pelechano laboratories for helpful scientific exchange. Computational analysis was performed on resources provided by the National Academic Infrastructure for Supercomputing in Sweden (NAISS), partially funded by the Swedish Research Council through grant agreement no. 2022-06725. This work was supported by the Initiative and Networking Fund of Helmholtz Young Investigators program of the Helmholtz Association (funding code: VH-NG-1535), by the Deutsche Forschungsgemeinschaft (DFG, grant agreement No 535634027) and by the European Research Council (ERC) under the European Union’s Horizon 2020 research and innovation program (grant agreement ERC-STG No 948544) to IP. IP also acknowledges funding from the Wallenberg Academy Fellow program (KAW 2023.0053), Stockholm University and Karolinska Institutet. VP acknowledges funding from an extension Wallenberg Academy Fellowship (KAW 2021.0167), the Swedish Research Council (VR 2020-01480 and 2024-03210) and Karolinska Institutet (SciLifeLab Fellowship, SFO, and KI funds). EKB acknowledges postdoctoral support from EMBO, the Swedish Society for Medical Research (SSMF) and KI.

## Contributions

E.F. performed and analysed all experiments; A.C.J. performed and analysed selected targeted MolMap experiments; E.F. and A.C.J performed bioinformatic analysis of the data. E.K.B helped designing, and analyzing MolMap assays. M.W. helped with data analysis. I.P. and E.F. wrote the manuscript with contributions from all authors. E.K.B. co-supervised part of the project. I.P. and V.P. conceived and supervised the project.

