## Supplementary material for "MolMap resolves functional chromatin responses to small-molecule perturbation": Fuerst_BioRxiv_2026_Extended

### Extended data

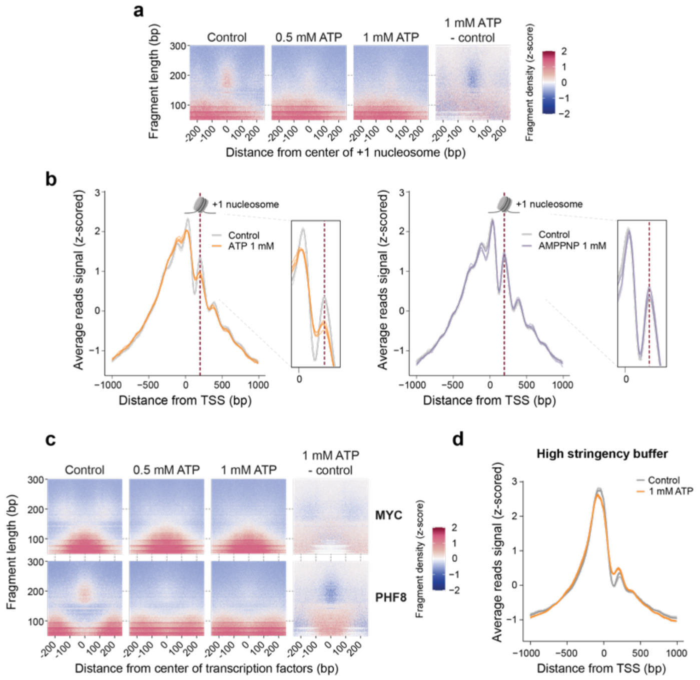


**Extended data Fig 1. MolMap captures chromatin structural remodeling at nucleosome and TF-associated sites.**

**a.** Fragment length vs. position heatmaps (V-plots) showing z-score–normalized MolMap fragment density around annotated +1 nucleosome showing the displacement of the nucleosome after ATP treatment in K562 nuclei. Coordinates are derived from the MNase-seq datasets from Kundaje et al., 2012[53]. The x-axis denotes the distance (±250 bp) from the center of binding sites, and the y-axis represents the length of MolMap fragments in base pairs. Heatmaps display fragment density across a ±250 bp window relative to nucleosome centers for control, 0.5 mM ATP, and 1 mM ATP-treated nuclei. The right panel shows the difference between 1 mM ATP and control signals. Fragment density is normalized per fragment length; red indicates enrichment, blue indicates depletion.

**b.** MolMap treatment with ATP and AMPPNP, a non-hydrolysable ATP analogue. Average MolMap read coverage profiles centered on transcription start sites (TSSs) ±1 kb in control vs 1 mM ATP (left panel, and control vs 1 mM AMPPNP (right panel). Signal tracks represent z-score–scaled coverage per 10 bp bin, with each line corresponding to a separate biological replicate. Dashed vertical red lines show the position of the +1 nucleosome. Insets show a magnified view of the +1 nucleosome region, highlighting the ATP-induced accessibility change at the +1 nucleosome region, which is absent with AMPPNP. These results indicate that ATP hydrolysis, rather than ATP binding alone, is required for the observed chromatin response.

**c.** V-plots showing z-score–normalized MolMap fragment density centered on genome-wide MYC and PHF8-associated sites in K562 nuclei, following treatment with control, 0.5 mM ATP, or 1 mM ATP. The x-axis denotes the distance (±250 bp) from the center of binding sites, and the y-axis represents the length of MolMap fragments in base pairs. Central short-fragment protection patterns are associated with MYC-bound regions; flanking patterns reflect nucleosome positioning. For PHF8 the nucleosome at 0 bp overlaps with PHF8-associated sites. The right panel shows differential signal between 1 mM ATP and control. Fragment density is normalized per fragment length; red indicates enrichment, blue indicates depletion.

**d.** Average MolMap read coverage profiles centered on transcription start sites (TSSs) ±1 kb in control and 1 mM ATP-treated K562 cells when a high stringency buffer with higher ionic strength was used. Signal tracks represent z-score–scaled coverage per 10 bp bin, with each line corresponding to a separate biological replicate.

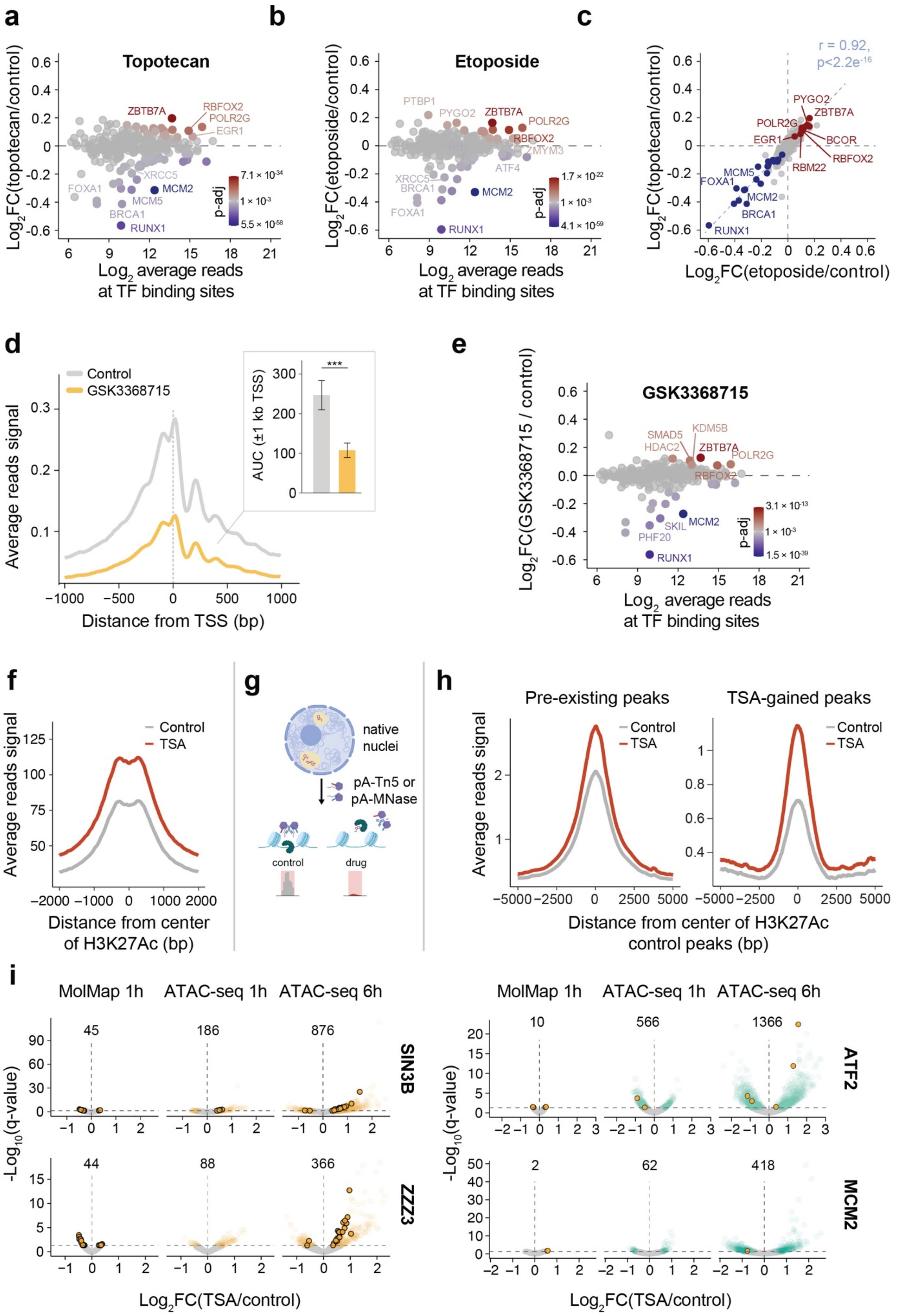


**Extended data Fig. 2**

**Extended data Fig 2: MolMap resolves chromatin responses to protein-targeting compounds**

**a.** MA plots of differential MolMap accessibility at aggregated transcription factor-associated sites following treatment of K562 nuclei with 2 µM topotecan. Each point represents one TF with colors indicating the direction and significance of accessibility changes (DESeq2 Wald test; adjusted $p < 0.001$, Benjamini–Hochberg correction), with red and blue showing increased and decreased accessibility.

**b.** MA plots of differential MolMap accessibility at aggregated transcription factor-associated sites following treatment of K562 nuclei with 2 µM etoposide. Each point represents one TF with colors indicating the direction and significance of accessibility changes (DESeq2 Wald test; adjusted $p < 0.001$, Benjamini–Hochberg correction), with red and blue showing increased and decreased accessibility.

**c.** Correlation of MolMap chromatin accessibility changes at TF-associated sites induced by topotecan and etoposide. Each point represents a TF, with $\log_2$ fold changes of aggregated TF-associated sites (drug vs. DMSO control) on the x- and y-axes. Points are colored by significance: increased (red) or decreased (blue) MolMap accessibility (DESeq2 Wald test; adjusted $p < 0.001$; Benjamini–Hochberg correction). A linear fit reveals high similarity in TF accessibility profiles between the two compounds. r pearson correlation coefficient = 0.92.

**d.** Average MolMap signal profiles at transcription start sites (TSSs) ±1 kb for control (DMSO) and 2 µM GSK3368715-treated nuclei. Signals represent normalized read coverage per 10 bp bin, averaged across all TSSs and merged across replicates for each condition. Right panel: average area under the curve (AUC) around 1 kb of the TSS (Statistical Significance: Two-sided Welch's t-test, ***$P < 0.001$). Read coverage per 10 bp bin is normalized to the internal *E. coli* DNA reference.

**e.** MA plot of MolMap accessibility changes at aggregated TF-associated sites following 2 µM GSK3368715 treatment. Each point represents one TF. Point color indicates the direction and significance of accessibility changes (DESeq2 Wald test; adjusted $p < 0.001$, Benjamini–Hochberg correction), with red and blue showing increased and decreased accessibility.

**f.** Average MolMap signal profiles at H3K27Ac sites ±2 kb for control and 20 µM trichostatin A-treated nuclei. Signals represent normalized read coverage per 10 bp bin, averaged across all H3K27Ac sites and merged across replicates for each condition. Read coverage per 10 bp bin is normalized to the internal *E. coli* DNA reference. TSA indicates trichostatin A.

**g.** Experimental set up of the targeted MolMap assays in native nuclei. Native nuclei isolation and subsequent treatment with ligand and an antibody against a target of interest: *i)* Tn5-based targeted MolMap: a protein A-Tn5 transposase fusion tagments the DNA in close proximity to the target. *ii)* MNase-based targeted MolMap: a protein A-MNase fusion cuts the DNA in close proximity to the target. pA = protein A.

**h.** Tn5-based targeted MolMap at H3K27Ac sites upon 20 µM trichostatin-A treatment. Average MolMap signal profiles at pre-existing H3K27Ac peaks ±5 kb identified in control samples (left) and new peaks quantified upon trichostatin A treatment (TSA-gained peaks, right). Signals represent library size-normalized read coverage per 10 bp bin, averaged across all H3K27ac peaks and merged across replicates for each condition. TSA indicates trichostatin A.

**i.** Volcano plots of accessibility changes at individual chromatin regulators sites (SIN3B, ZZZ3, ATF2 and MCM2) in MolMap 1h, ATAC-seq 1h and ATAC-seq 6h upon 20 µM of trichostatin A treatment. Each point represents a TFBS. Numbers above each plot indicate the number of significant TFBSs detected in the corresponding dataset (DESeq2 Wald test, adjusted $p < 0.05$, Benjamini–Hochberg correction). Orange points with black contour: pinpoint TFBSs with a significant change in MolMap signal. TSA indicates trichostatin A.

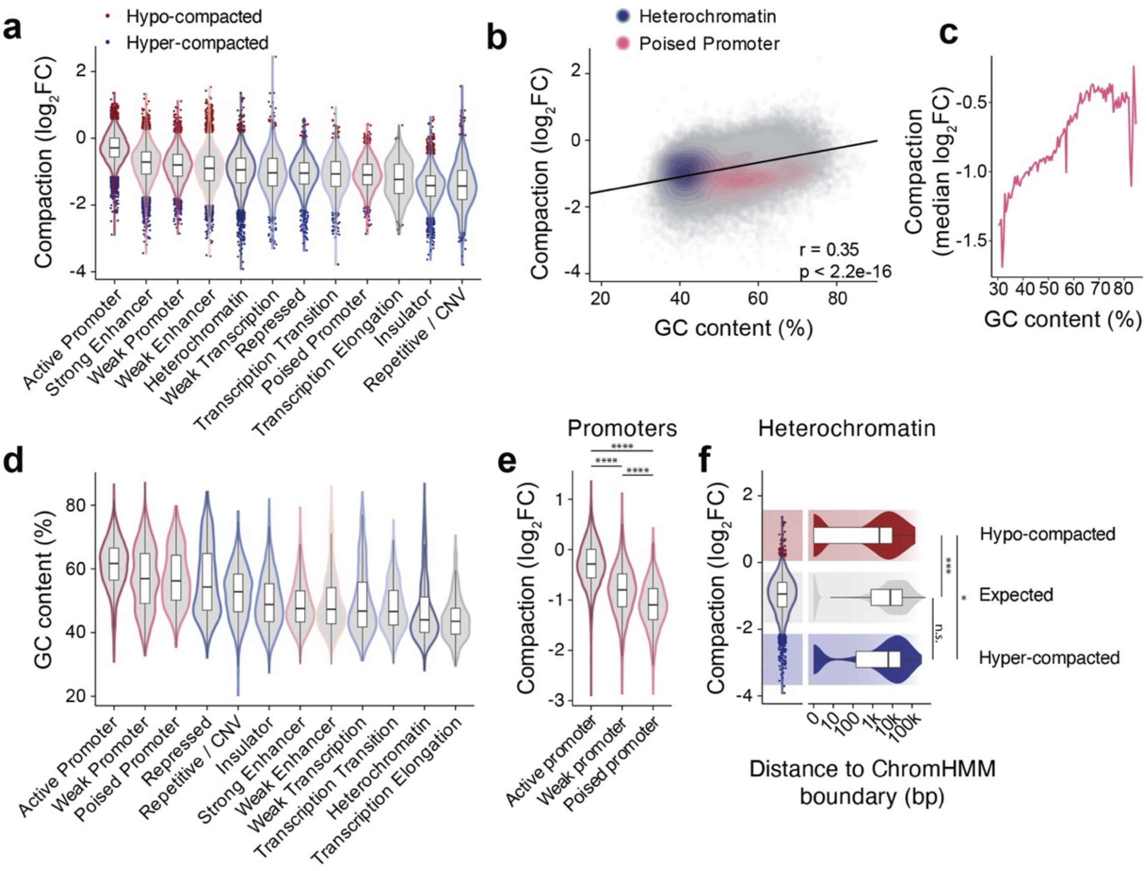


**Extended data Fig 3: Polyamine-induced chromatin compaction is resolved by MolMap in native nuclei**

Analysis of $\log_2$ MolMap chromatin accessibility fold changes upon 5 mM spermidine treatment. For this analysis, chromatin compaction is operationally defined as the reduction in MolMap accessibility following spermidine treatment. In this context we define compaction $\log_2$ fold changes ($\log_2FC$) as the equivalent $\log_2$ MolMap chromatin accessibility fold change with the sign change.

**a.** Distribution of peak $\log_2$ fold changes across ChromHMM-defined chromatin states upon spermidine treatment (5 mM). Regions classified as hyper-compacted (blue) (z-score < -2) or hypo-compacted (red) (z-score > 2) are highlighted relative to the background distribution within each chromatin state. Box plots: horizontal lines define the median and boxes the 25$^{th}$ and 75$^{th}$ percentiles; whiskers 1.5 × the interquartile range.

**b.** Correlation between chromatin compaction and GC content. Each point represents an individual genomic locus. Red line: fitted regression trend estimated using a linear regression model (Pearson's correlation *r* = 0.35).

**c.** Median chromatin compaction across bins of GC content. Regions were grouped by GC percentage, and the median compaction ($\log_2FC$) was calculated for each bin: bin width: 0.5% GC content.

**d.** Distribution of GC content across spermidine significant peaks grouped according to their respective ChromHMM chromatin states. Box plots: horizontal lines define the median and boxes the 25$^{th}$ and 75$^{th}$ percentiles; whiskers represent 1.5 × the interquartile range.

**e.** Compaction changes ($\log_2FC$) at promoter-associated chromatin states. Violin plots show compaction distributions for active, weak, and poised promoters. Wilcoxon test; ****: $p < 0.0001$.

**f.** Distance of hypo-compacted (z-score > 2), hyper-compacted (z-score < -2) heterochromatin regions and heterochromatin regions exhibiting expected ($-2 \leq$ z-score $\leq 2$) compaction behavior to the nearest ChromHMM state boundary (bp = base pairs). Violin plots show the distribution of distances (bp), Wilcoxon test; ***: $p < 0.001$, *: $p < 0.05$. Box plots: horizontal lines define the median and boxes the 25$^{th}$ and 75$^{th}$ percentiles; whiskers represent 1.5 × the interquartile range.

**Extended data Fig 4.**

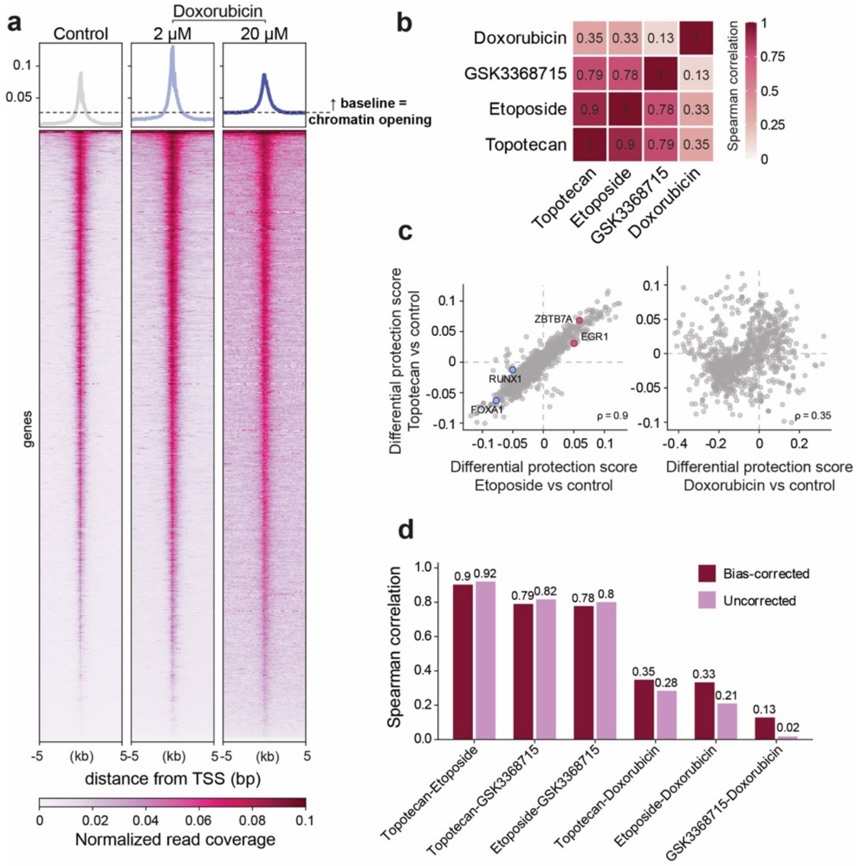


**Extended data Fig 4: MolMap resolves compound- and mechanism-dependent chromatin-responses**

**a.** MolMap reveals global chromatin remodeling at at transcription start sites following doxorubicin treatment. Average MolMap read coverage profiles centered on transcription start sites (TSSs) ±5 kb in control, 2 µM, and 20 µM doxorubicin-treated nuclei. Signal is normalized to the internal *E. coli* DNA reference and averaged across refGene-annotated TSSs. The heatmaps below show coverage across all TSSs sorted by decreasing accessibility in the control condition. The elevated baseline under 20 µM doxorubicin reflects genome-wide chromatin opening.

**b.** Pairwise correlations of Tn5-bias-corrected motif-associated differential protection scores following treatment with topotecan, etoposide, GSK3368715 and doxorubicin. Scores quantify treatment-induced changes in local Tn5 protection around transcription-factor motifs relative to the corresponding matched control. Spearman correlation coefficients are shown. The related topoisomerase inhibitors topotecan and etoposide show the highest concordance, whereas doxorubicin shows a clearly distinct response pattern.

**c.** Representative pairwise comparisons of motif-associated differential protection scores. Topotecan and etoposide show highly concordant responses, whereas topotecan and doxorubicin show substantially lower concordance. Each point represents one transcription-factor motif from the JASPAR CORE vertebrate motif collection[52]. Labelled motifs indicate transcription factors that are also highlighted by the aggregated TF-associated site analysis shown in Extended Data Fig. 2c.

**d.** Effect of Tn5-bias correction on pairwise compound comparisons. Correlations between motif-associated differential protection scores were calculated before and after correction for sequence-dependent Tn5 insertion bias. The relative relationships among compounds are preserved after correction, indicating that the observed compound-dependent response patterns are not driven by Tn5 sequence preference.

The differential protection scores in panels b-d are calculated using the algorithm TOBIAS[38]. TOBIAS differential binding scores are here interpreted as changes in local chromatin protection around transcription-factor motifs, rather than as direct measurements of transcription-factor occupancy.

**Extended data Fig 5.**

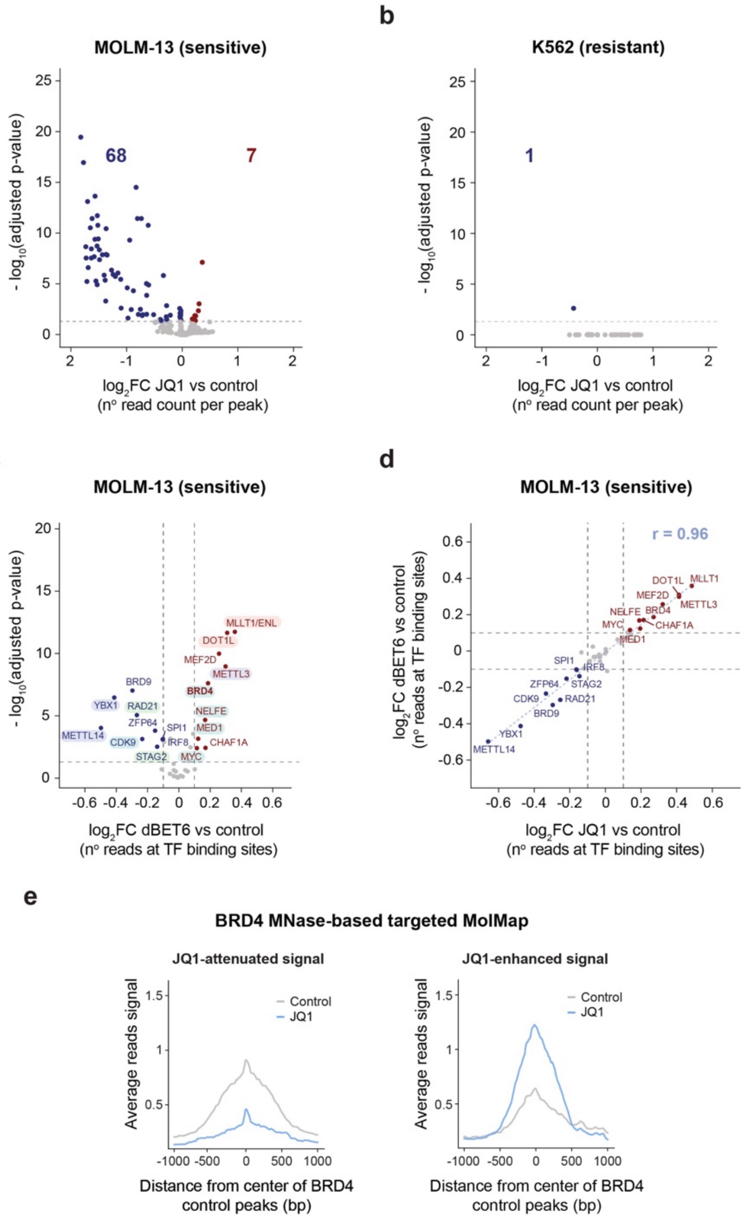


**Extended data Fig 5: MolMap distinguishes sensitive and resistant chromatin responses to BRD4 inhibitors in leukemia cells**

**a–b.** Volcano plots of peak-level chromatin accessibility changes after 20 µM JQ1 treatment in MOLM-13 (**a**) or K562 (**b**). Significant peaks (DESeq2 Wald test, adjusted $p < 0.05$; Benjamini–Hochberg correction) are highlighted in red (up) and blue (down).

**c.** Volcano plot showing MolMap chromatin accessibility changes at aggregated transcription factor (TF) -associated sites in MOLM-13 cells treated with 20 µM dBET6 or DMSO. Red: increased accessibility; blue: decreased (DESeq2 Wald test, adjusted $p < 0.05$, $|\log_2 FC| > 0.1$). The analysis shows strong overlap with the effect of JQ1. Hits are grouped by known functional relationship to BRD4 biology and divided in classes as in Figure 5.

**d.** Correlation of MolMap fold changes across TF-associated sites after JQ1 vs. dBET6 treatment at 20 µM, indicating highly concordant BRD4-linked chromatin responses. *r* Pearson correlation coefficient = 0.96

**e.** BRD4 MNase-based targeted MolMap in MOLM-13 cells. Average signal profiles centred on BRD4 peaks (±1 kb) for control (DMSO) and JQ1 treated nuclei (20 µM) of MOLM-13 cells. Left: JQ1-attenuated BRD4-associated signal: pre-existing BRD4 peaks identified in control samples that lose MolMap signal upon JQ1 exposure. Right: JQ1-enhanced response detected upon JQ1 treatment. Signals represent CPM-normalized MolMap read coverage averaged across all peaks of the indicated genomic window relative to peak centers.
